# Multimodal Magnetic Resonance Imaging Biomarkers of Pediatric Traumatic Brain Injury Identified by Ensemble Learning Are Associated with Psychopathology

**DOI:** 10.64898/2026.09.01.748387

**Authors:** Elizabeth Martin, Kai Wu, Xiaobo Li

**Author notes:** Corresponding authors: Xiaobo Li, PhD Department of Biomedical Engineering New Jersey Institute of Technology New Jersey, USA Email Address Elizabeth Martin, PhD Department of Biomedical Engineering New Jersey Institute of Technology New Jersey, USA.

## Abstract

**Background:** Pediatric traumatic brain injury (TBI) is associated with increased psychopathology, but the neural alterations underlying this vulnerability remain poorly understood. We used interpretable machine learning to identify multimodal MRI features associated with pediatric TBI and their relationship with psychopathology.

**Methods:** Baseline multimodal MRI data from 775 children aged 9-10 years in the Adolescent Brain Cognitive Development Study were analyzed, including 365 children with TBI and 410 controls. Structural MRI, diffusion MRI, and resting-state functional MRI features were evaluated using nested cross-validated ensemble classification. Permutation-based feature importance identified discriminative features, which were subsequently examined using partial least squares (PLS) regression in the TBI and control groups.

**Results:** Classification performance was above chance across models, with accuracy ranging from 0.612-0.665 and AUC from 0.644-0.714, and was comparable in an independent validation sample. Twenty neuroimaging features spanning all three modalities were reliably discriminative, including measures of white matter microstructure, gray matter volume, regional functional variability, and functional connectivity. In the TBI group, a significant PLS component was associated with overall psychopathology scores (r = 0.22, permuted p = 0.008), whereas no significant component was identified in controls. Ten discriminative features contributed substantially to this association.

**Conclusions:** Pediatric TBI is characterized by a distributed multimodal neuroimaging signature specifically associated with psychopathology following injury. Interpretable machine learning combined with multivariate brain-behavior modelling may help identify neural substrates of psychiatric vulnerability after pediatric TBI.

## INTRODUCTION

Traumatic brain injury (TBI) is a leading cause of death and acquired disability in children and adolescents worldwide.^1^ Because the pediatric brain is undergoing rapid structural and functional maturation, injuries sustained during this critical developmental period may disrupt normative neurodevelopmental trajectories, with consequences extending well beyond the acute injury phase.^2–4^

Among the most clinically significant sequelae of pediatric TBI is an increased risk of psychopathology. Novel psychiatric disorders, including secondary attention-deficit/hyperactivity disorder, personality change, and internalizing and externalizing disorders, occur more frequently following TBI, even in children without a prior psychiatric history.^4, 5^ Prospective studies have demonstrated that emotional and behavioral difficulties can persist for months or years after injury, with even mild TBI associated with an increased risk of affective and behavioral problems.^6–9^ Despite this well-established relationship, the neurobiological mechanisms through which TBI increases vulnerability to psychopathology remain incompletely understood, in part because of the limitations of traditional univariate analyses in identifying distributed neural alterations.^10, 11^

Magnetic resonance imaging (MRI) provides complementary measures of TBI-related neuropathology. Structural MRI (sMRI) detects TB-related alterations in gray matter morphology, diffusion MRI (dMRI) characterizes white matter microstructural integrity, and resting-state functional MRI (rs-fMRI) captures changes in brain function and network organization.^12–28^ These modalities reflect distinct aspects of brain injury pathology, and multimodal approaches have demonstrated improved characterization of TBI compared with single-modality analyses.^10, 29^

The Adolescent Brain Cognitive Development (ABCD) Study provides an unprecedented opportunity to investigate these relationships in a large, population-based sample. The ABCD Study enrolled over 11,500 children aged 9-10 years from 21 sites across the United States, with approximately 3% reporting a history of TBI.^30–32^ Its comprehensive multimodal neuroimaging protocol, combined with detailed assessments of mental health, enables investigation of brain-behavior relationships at a scale not previously possible.^30, 33^ Previous analyses of the ABCD cohort have identified associations between pediatric TBI, altered brain structure and function, and increased psychopathology, but no study has comprehensively examined multimodal MRI features and their relationship with psychopathology using an interpretable machine learning framework.^7, 31, 34^

Machine learning methods are well suited to multimodal neuroimaging because they can model complex, high-dimensional relationships that are difficult to detect using conventional statistical approaches. Ensemble learning further improves predictive performance by combining multiple complementary classifiers, often producing more robust and generalizable models than individual algorithms. However, predictive performance alone provides limited biological insight, as ensemble models are frequently criticized for functioning as “black boxes”.^35–37^ Permutation-based feature importance offers one solution by identifying the imaging features that contribute most consistently to classification.^38^ These discriminative features can then be examined using multivariate approaches such as partial least squares (PLS) regression to determine how patterns of brain alteration relate to clinical outcomes.^39, 40^ This combination of interpretable machine learning and multivariate brain-behavior modelling provides a framework that extends beyond classification toward biological interpretation.

The present study addressed three aims using baseline data from the ABCD Study. First, we applied a nested cross-validated ensemble learning framework to determine whether multimodal MRI features could distinguish children with a history of TBI from matched controls. Second, permutation-based feature importance was used to identify the structural and functional neuroimaging features that contributed most consistently to classification. Third, these discriminative features were entered into a PLS regression to examine their multivariate associations with dimensions of psychopathology. Together, this pipeline provides a data-driven approach for identifying the neural correlates of pediatric TBI and characterizing their relationship with psychopathology. We hypothesized that (1) ensemble learning would successfully classify children with TBI from controls using multimodal MRI features, (2) discriminative features would span structural, diffusion, and functional imaging modalities, and (3) these features would be significantly associated with psychopathology.

## MATERIALS AND METHODS

### Participants

The present study analyzed de-identified baseline data from the Adolescent Brain Cognitive Development (ABCD) Study (Release 6.0), a nationwide cohort that enrolled 11,875 US children aged 9-10 years across 21 sites through school-based, demographically stratified recruitment. The ABCD baseline data release includes multimodal neuroimaging data and extensive demographic and clinical data. The ABCD study was approved by the institutional review board (IRB) of the University of California, San Diego and each data collection site. Informed consent and informed assent were obtained from parents and participants, respectively. The current study is a secondary analysis of de-identified data and therefore IRB approval was waived.

### TBI group definition

Parents/guardians of all ABCD Study participants completed the Ohio State University TBI Identification Method.^41^ Using the Ohio State method, participants are categorized into six categories: no TBI (no head/neck injury), improbable TBI (no TBI, or TBI without loss-of-consciousness or memory loss), possible mild TBI (no loss-of-consciousness, but with memory loss), mild TBI (loss-of-consciousness lasting less than 30 minutes), moderate TBI (loss-of-consciousness lasting 30 minutes to 24 hours), and severe TBI (loss-of-consciousness lasting longer than 24 hours). Participants were considered for the TBI subject pool if parental reports characterized them as having at least possible mild TBI. Four hundred and forty-four participants with TBI were identified using these criteria. A pool of control subjects was identified as participants with no head or neck injury/impact, or improbable TBI, characterized as no TBI or TBI with no memory loss or loss-of-consciousness. To both subject groups, the following exclusion criteria were then applied: no schizophrenia, autism spectrum disorder, no recorded use of non-stimulant psychotropic medications. The ABCD study’s imaging data quality metric was then applied for each separate imaging modality, and any subject with unacceptable data for any imaging modality was excluded. Following these exclusion criteria, the final number of TBI subjects included in analysis was 365 and the final number of control subject included in analysis was 410.

### ABCD Study Neuroimaging Data

Baseline neuroimaging data were obtained from the tabulated imaging release of the Adolescent Brain Cognitive Development (ABCD) Study (Release 6.0). The ABCD Study acquired multimodal MRI data from participants at 21 sites across the United States using harmonized 3T scanner protocols from three manufacturers (Siemens, GE, and Philips). MRI acquisition and preprocessing followed the standardized ABCD imaging pipeline as previously described.

Images were acquired on harmonized 3T Siemens, GE and Philips scanners. Scanner manufacturer, scanner ID and software version were included as covariates to account for site-related effects.^30, 42, 43^

#### Diffusion tensor imaging

ROI-averaged fractional anisotropy (FA) and mean diffusivity (MD) were extracted from 42 white matter tracts defined by the AtlasTrack atlas. FA reflects the directional coherence of water diffusion along axonal fibers and is sensitive to alterations in axonal organization and myelination, whereas MD reflects the overall magnitude of water diffusion and can increase with tissue disruption. FA and MD were therefore included as complementary measures of white matter microstructural integrity.^30, 42^

#### Structural MRI

Gray matter morphology was characterized from T1-weighted structural MRI. ROI-averaged total volume and thickness were extracted for cortical regions defined by the Destrieux atlas and for subcortical structures defined by the FreeSurfer subcortical segmentation (aseg atlas).^44^ Fifteen subcortical and 148 cortical ROIs were included in the feature pool, providing measures of regional gray matter morphology across cortical and subcortical structures.

#### Resting-state functional MRI

Resting-state fMRI features comprised (1) regional BOLD temporal variance across 148 Destrieux cortical ROIs and (2) 169 within- and between-network functional connectivity measures derived from the Gordon parcellation. Temporal variance was used to characterize regional fluctuations in spontaneous BOLD activity, while functional connectivity measures captured the strength of functional coupling within and between canonical brain networks.^42, 44 45 46^

### Statistical Analysis

Analyses were conducted in three stages: feature selection, ensemble classification, and PLS regression, each described in detail below. Figure 1 presents the analytic pipeline. All analyses were performed in Python (version 3.12.1) using the scikit-learn machine learning library. The outcome variable (TBI/Control) was binary-encoded prior to modeling.

**Figure 1.**
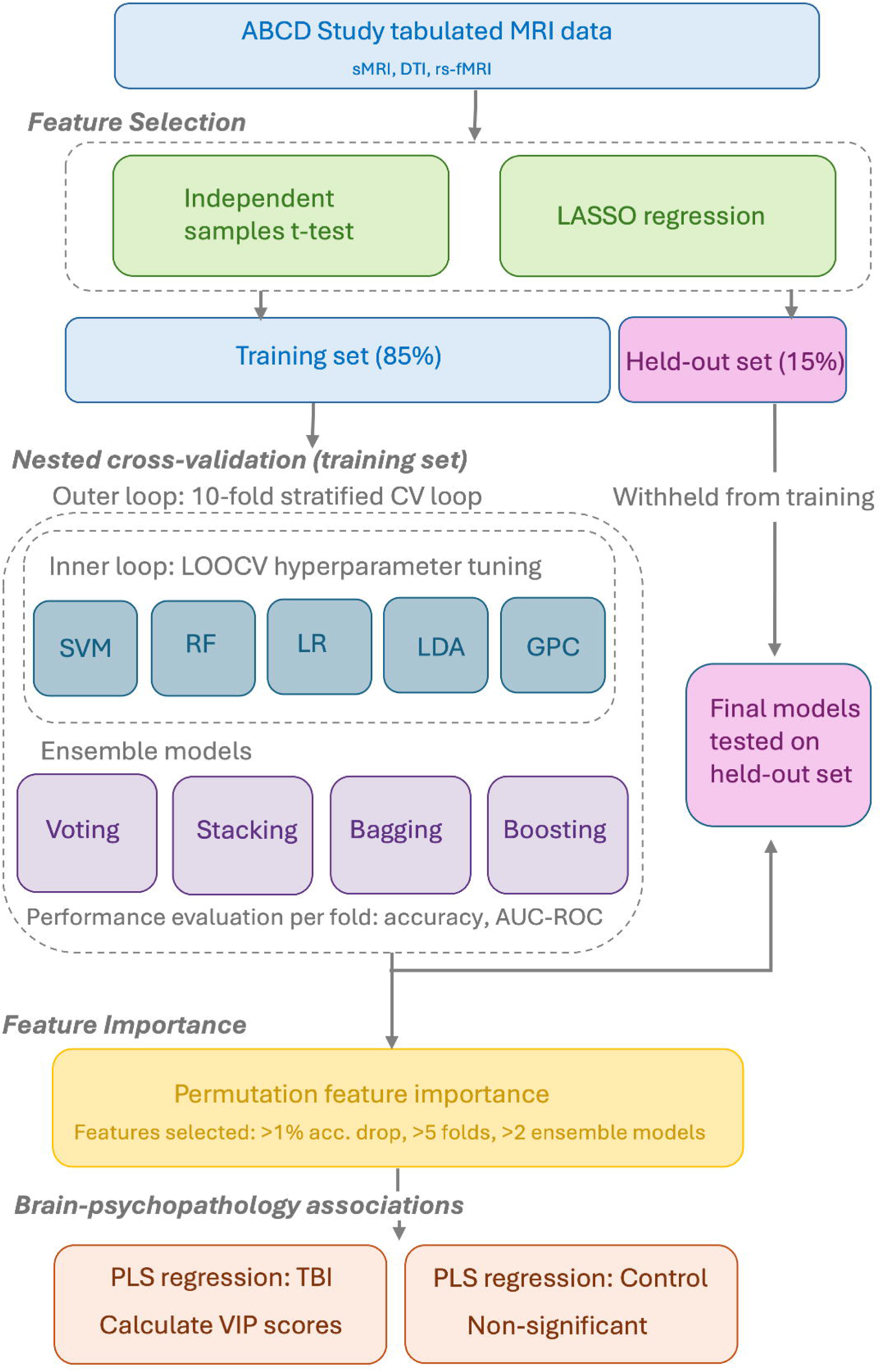
Analysis pipeline for ensemble classification and PLS regression. Multimodal MRI features (DTI, sMRI, rs-fMRI) from the ABCD Study underwent two-stage feature selection (independent samples t-tests, LASSO regression), retaining combined features from both. Data were divided into a development set (85%) and an independent holdout set (15%), stratified by group. Within the development set, five base classifiers (SVM, RF, LR, LDA, GPC) and four ensemble methods (soft voting, stacking, bagging, gradient boosting) were evaluated using nested cross-validation (outer: 10-fold stratified CV; inner: LOOCV with grid search hyperparameter tuning). Final models were applied to the holdout set to assess generalization. Permutation-based feature importance identified features producing >1% accuracy reduction in ≥5 outer folds across ≥2 ensemble models. Retained features were entered into separate PLS regression models for TBI and control groups, with CBCL syndrome scores as outcomes. Component significance was assessed via permutation testing (1,000 permutations); features with VIP scores >1.0 were identified as principal brain-psychopathology drivers. Abbreviations: DTI = diffusion tensor imaging; sMRI = structural MRI; rs-fMRI = resting-state functional MRI; SVM = support vector machine; RF = random forest; LR = logistic regression; LDA = linear discriminant analysis; GPC = Gaussian process classifier; LOOCV = leave-one-out cross-validation; PLS = partial least squares; CBCL = Child Behavior Checklist; VIP = variable importance in projection.

#### Model development and evaluation framework

Feature selection was performed on the original feature pool of 685 combined features using two-sample t-test and lasso regression. The 40 features with the lowest p-value in the t-test, and the 40 features with the highest absolute lasso coefficient were combined, resulting in 67 final features after removal of overlapping features. This contained features of all modalities, although within sMRI features, only volume features met the selection criteria.

Classification performance was evaluated using a nested cross-validation framework to obtain unbiased estimates of generalization error while simultaneously optimizing hyperparameters. The outer loop employed 10-fold stratified cross-validation, preserving class proportions across folds, to assess model performance on held-out data. The inner loop used Leave-One-Out cross-validation (LOOCV) for hyperparameter tuning via grid search, optimizing for classification accuracy. All feature imputation (mean substitution for missing values) and scaling (z-score standardization, where applicable) were performed within each training fold to prevent data leakage. Prior to model training, 15% of the sample was withheld as an independent validation set using stratified random sampling. The remaining 85% of the sample was used for all subsequent model development steps. The holdout set was not used or inspected during nested cross-validation, and was only accessed once a final trained model was available.

#### Base classifiers

Five base classifiers were evaluated: (1) Support Vector Machine (SVM) with radial basis function kernel, with regularization parameter C and kernel width γ optimized via grid search; (2) Random Forest (RF), with 300 decision trees with maximum depth, minimum leaf samples, and feature subset size tuned via grid search; (3) Logistic Regression (LR) with L2 regularization, with the regularization strength C optimized via grid search; (4) Linear Discriminant Analysis (LDA), fit without additional hyperparameter tuning; and (5) Gaussian Process Classifier (GPC) with RBF kernel, with kernel length scale optimized via grid search. For each outer fold, each base classifier was independently tuned on the inner training data and evaluated on the outer held-out test set. Performance metrics calculated per-fold were accuracy, and area under the receiver operating characteristic curve (AUC-ROC), whereby a perfect classifier would result in an AUC-ROC of 1.0, and a classifier performing worse than random chance would result in an AUC-ROC of <0.5.

#### Ensemble methods

Four ensemble strategies were constructed within each outer fold using the tuned base classifiers as components. A soft-voting ensemble combined the probability outputs of all five tuned base classifiers. A stacking ensemble used the five tuned base classifiers as first-level learners and a logistic regression model as the meta-learner, trained using 5-fold cross-validation on the training fold. A bagging ensemble was constructed by applying Bootstrap Aggregating (BaggingClassifier; 50 estimators, 80% bootstrap sample size) independently to each of the five tuned base classifiers, and predictions were averaged across all resulting models. Finally, a gradient boosting classifier (GradientBoostingClassifier with default hyperparameters) was trained directly on the training fold. Classification accuracy and AUC-ROC were computed for each ensemble on the outer held-out test fold, to calculate mean performance metrics across all 10 folds.

#### Feature importance

Permutation-based feature importance was computed (permutation_importance) for the voting, stacking, and gradient boosting ensembles across all 10 outer cross-validation folds (10 permutation repeats per fold, random state fixed for reproducibility) using and averaged across folds.^38^ Features were retained for downstream analysis if they met the following criteria: a mean reduction in accuracy of greater than 1% upon permutation in at least five of the ten outer folds, replicated across at least two of the three ensemble models.

#### PLS Regression

Psychopathology was assessed using the caregiver-completed Child Behavior Checklist (CBCL). Eight syndrome scales (Anxious/Depressed, Withdrawn/Depressed, Somatic Complaints, Social Problems, Thought Problems, Attention Problems, Rule-Breaking Behavior, and Aggressive Behavior). ^47^ The eight-syndrome structure has been validated cross-nationally across 30 societies in a confirmatory factor analysis of over 58,000 children aged 6 to 18.^33, 48^ All eight syndrome scale scores were entered as outcome variables in the PLS regression analyses.

PLS regression was used to examine the multivariate association between the selected neuroimaging features and CBCL syndrome scores.^40, 49^ Separate PLS models were estimated for the TBI group and the control group, and permutation testing (1,000 permutations) was used to assess the significance of each extracted component. Variable importance in projection (VIP) scores exceeding 1.0 were used to identify the features most strongly driving each significant component.^50^

## RESULTS

### Sample Characteristics

The final analytic sample comprised 365 children with a history of TBI and 410 typically developing controls drawn from the ABCD Study. Groups did not differ significantly on sex, age, socioeconomic status (household income) or ethno-racial identity. Of the 365 children with TBI, 261 had possible mild TBI, 99 had mild TBI, 3 had moderate TBI, and 2 had severe TBI. Statistical comparisons of sample characteristics are shown in **Table 1**.

**Table 1.** Participant characteristics. TBI = participants with traumatic brain injury; LOC = loss-of-consciousness; mem. loss = memory loss; p = p-value for independent; t-test for continuous variables, and for chi-square test for categorical variables

|  | TBI (n = 365) |  | Control (n = 410) |  | p |
| --- | --- | --- | --- | --- | --- |
|  | mean | (sd) | mean | (sd) |  |
| **Age (years)** | 10.1 | 0.62 | 10.1 | 0.63 | 0.89 |
|  | **n** | **(%)** | **n** | **(%)** |  |
| **Sex** |  |  |  |  |  |
| Male | 219 | (60.0) | 217 | (52.9) | 0.056 |
| Female | 146 | (40.0) | 193 | (47.1) |  |
| **Household Income** |  |  |  |  | 0.094 |
| < 50k | 65 | (17.8) | 84 | (20.5) |  |
| 50k to 100k | 96 | (26.3) | 126 | (30.7) |  |
| > 100k | 175 | (47.9) | 185 | (45.1) |  |
| Don't know | 16 | (4.4) | 7 | (1.7) |  |
| Decline to answer | 11 | (3.0) | 8 | (1.9) |  |
| **Ethno-racial identity** |  |  |  |  | 0.12 |
| Hispanic | 58 | (15.8) | 69 | (16.8) |  |
| White (non-Hispanic) | 221 | (60.5) | 275 | (67.1) |  |
| Black (non-Hispanic) | 39 | (10.7) | 23 | (5.6) |  |
| Asian/Pacific Islander (non-Hispanic) | 4 | (1.1) | 7 | (1.7) |  |
| Multiracial (Hispanic) | 4 | (1.1) | 4 | (0.9) |  |
| Multiracial (non-Hispanic) | 36 | (9.9) | 29 | (7.1) |  |
| Other (non-Hispanic) | 2 | (0.5) | 3 | (0.7) |  |
| **TBI Severity** |  |  |  |  | <0.001 |
| No head or neck injury/impact | 0 | (0.0) | 326 | 79.5 |  |
| Improbable TBI (no TBI or TBI w/o LOC or mem. loss) | 0 | (0.0) | 84 | 20.5 |  |
| Possible mild TBI (TBI w/o LOC but mem. loss) | 261 | (70.1) | 0 | (0.0) |  |
| Mild TBI (TBI w/LOC < 30 min) | 99 | (26.8) | 0 | (0.0) |  |
| Moderate TBI (TBI w/LOC 30 min - 24 hrs) | 3 | (0.8) | 0 | (0.0) |  |
| Severe TBI (TBI w/ LOC > 24 hrs) | 2 | (0.5) | 0 | (0.0) |  |

### Base Classifier Performance

Performance exceeded chance across all base classifiers (Table 2). The SVM achieved the highest mean accuracy (ACC = 0.663, AUC = 0.710), while RF showed the lowest overall performance (ACC = 0.623, AUC = 0.663). Performance of LR, LDA and GPC was comparable, with AUC values consistently exceeding classification accuracy.

**Table 2.**
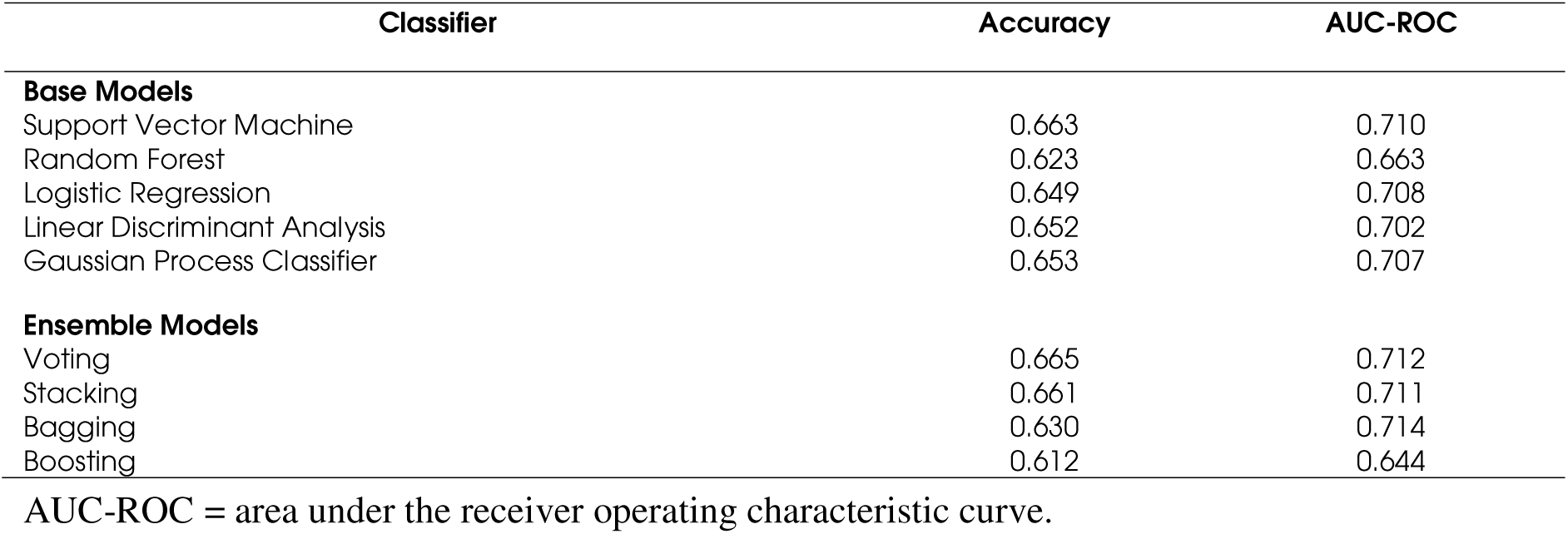
Mean classification accuracy and AUC-ROC for each classifier across 10 outer cross-validation folds.

| Classifier | Accuracy | AUC-ROC |
| --- | --- | --- |
| **Base Models** |  |  |
| Support Vector Machine | 0.663 | 0.710 |
| Random Forest | 0.623 | 0.663 |
| Logistic Regression | 0.649 | 0.708 |
| Linear Discriminant Analysis | 0.652 | 0.702 |
| Gaussian Process Classifier | 0.653 | 0.707 |
| **Ensemble Models** |  |  |
| Voting | 0.665 | 0.712 |
| Stacking | 0.661 | 0.711 |
| Bagging | 0.630 | 0.714 |
| Boosting | 0.612 | 0.644 |

AUC-ROC = area under the receiver operating characteristic curve.

### Ensemble Classifier Performance

Ensemble performance was broadly comparable to that of the best-performing base classifiers (Table 2). The soft voting ensemble achieved the highest classification accuracy (ACC = 0.665, AUC = 0.712), while bagging produced the highest AUC (0.714). Gradient boosting showed the weakest performance among the ensemble approaches.

### Independent Sample Validation

Base classifier performance on the holdout sample was broadly consistent with cross-validated estimates, with no evidence of substantial overfitting. The SVM again achieved the highest accuracy among base classifiers (ACC = 0.659, AUC = 0.687), closely followed by logistic regression and the GPC (both ACC = 0.650; AUC = 0.673 and 0.693 respectively). The GPC achieved the highest AUC among base classifiers on the holdout sample, marginally exceeding the SVM. LDA and RF produced the lowest holdout accuracy (both ACC = 0.626), with RF also showing the lowest AUC among base classifiers (0.620). The rank ordering of classifiers on the holdout sample was largely preserved relative to the cross-validated results, supporting the stability of the models.

Ensemble performance on the holdout sample was similarly consistent with cross-validated estimates. The soft voting and stacking ensembles again achieved the highest accuracy (both ACC = 0.650), with stacking achieving a marginally higher AUC (0.675 vs 0.673). Bagging achieved the highest AUC among ensemble methods on the holdout sample (AUC = 0.680), consistent with its pattern across the cross-validation folds. Gradient boosting produced the lowest accuracy and AUC on the holdout sample (ACC = 0.585, AUC = 0.614). Across both base classifiers and ensemble methods, holdout accuracy ranged from 0.585 to 0.659 and AUC ranged from 0.614 to 0.693.

### Permutation Feature Importance

Twenty neuroimaging features satisfied the permutation importance criterion (Table 3), spanning all three imaging modalities. Discriminative DTI features comprised reduced FA in the left fornix and right striatal-inferior frontal tract and elevated MD in the forceps major. Structural MRI features included gray matter volumes in the left central sulcus, left pallidum, left precuneus, bilateral occipital regions, and the right calcarine sulcus. The largest group of discriminative features derived from rs-fMRI, including regional temporal variance across frontal, parietal, occipital, temporal, insular and cingulate cortices, together with cingulo-parietal functional connectivity. The 20 selected features were subsequently carried forward as predictors in the PLS regression analyses (Figure 2).

**Figure 2.**
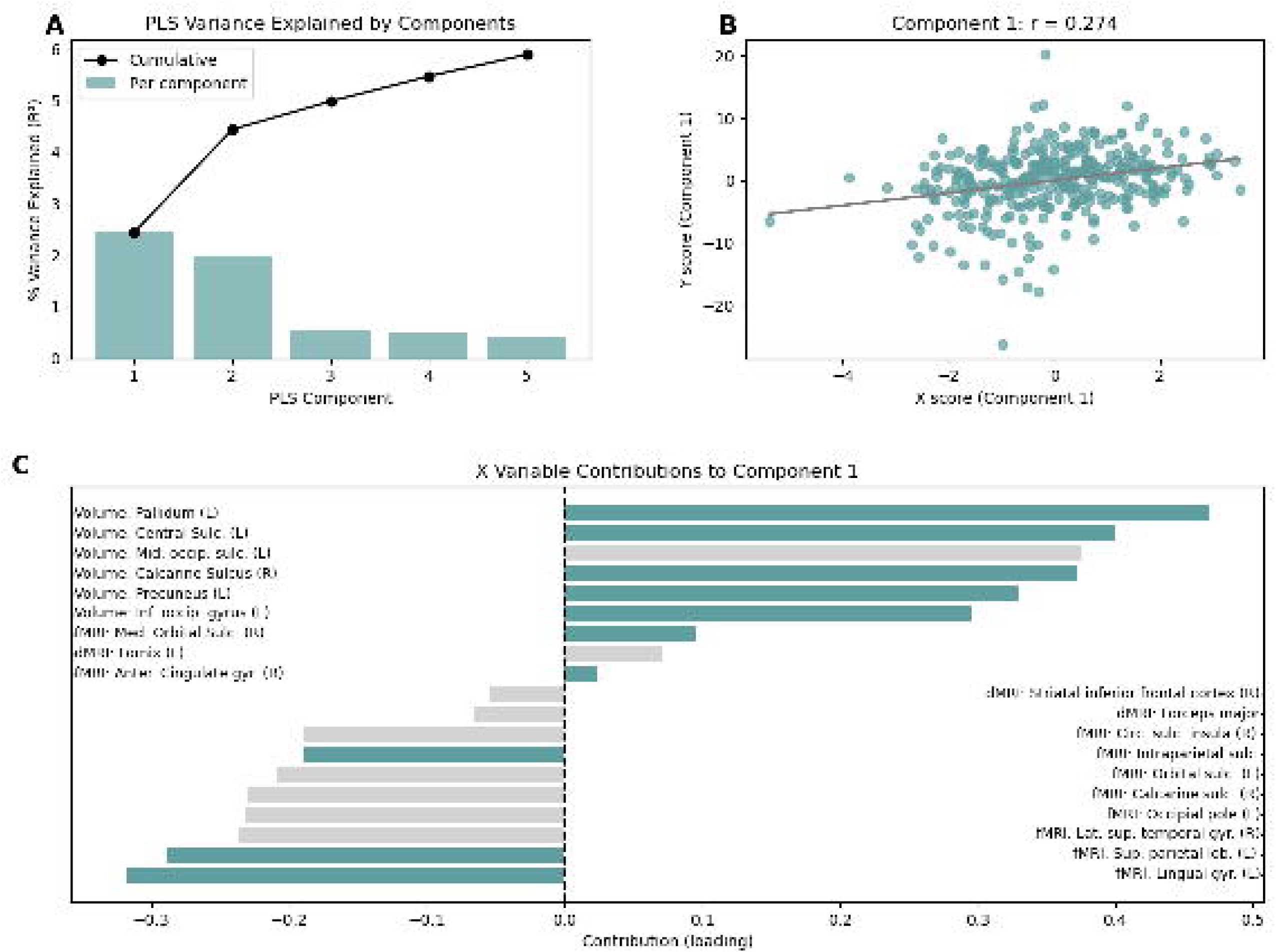
Sankey diagram showing the functional/anatomical categorization of important features in the classification and PLS models. Column one shows the 20 features meeting the criteria for feature importance in the ensemble classification models, categorized by region in the first column, imaging modality in the second column, and PLS VIP score in the third column.

**Table 3.** Neuroimaging features meeting the permutation importance criterion (>1% accuracy reduction in ≥5 outer folds across ≥2 ensemble models).

| Feature | Modality | Hemisphere | PLS VIP > 1 |
| --- | --- | --- | --- |
| FA: fornix | DTI | Left |  |
| FA: striatal inferior frontal cortex | DTI | Right |  |
| MD: forceps major | DTI | Bilateral |  |
| Volume: central sulcus | sMRI | Left | ☑ |
| Volume: calcarine sulcus | sMRI | Right | ☑ |
| Volume: pallidum (subcortical) | sMRI | Left | ☑ |
| Volume: middle occipital sulcus & lunatus sulcus | sMRI | Left |  |
| Volume: precuneus | sMRI | Left | ☑ |
| Volume: inferior occipital gyrus & sulcus | sMRI | Left | ☑ |
| Temporal variance: orbital sulci | fMRI | Left |  |
| Temporal variance: calcarine sulcus | fMRI | Right |  |
| Temporal variance: medial orbital sulcus | fMRI | Right | ☑ |
| Temporal variance: circular sulcus of insula, inf. segment | fMRI | Right |  |
| Temporal variance: superior temporal gyrus, lateral aspect | fMRI | Right |  |
| Temporal variance: intraparietal & transverse parietal sulci | fMRI | Left | ☑ |
| Temporal variance: superior parietal lobule | fMRI | Left | ☑ |
| Temporal variance: lingual gyrus | fMRI | Left |  |
| Temporal variance: anterior cingulate gyrus & sulcus | fMRI | Right | ☑ |
| Temporal variance: occipital pole | fMRI | Left |  |
| FC: cingulo-parietal-cingulo-parietal | fMRI | Bilateral |  |

FA = fractional anisotropy; MD = mean diffusivity; DTI = diffusion tensor imaging; sMRI = structural MRI; fMRI = resting-state functional MRI; rs-fMRI = resting-state fMRI; FC = functional connectivity. ☑ indicates VIP score > 1 in the TBI-group PLS regression, indicating important contribution to the regression.

### Partial Least Squares Regression: Brain-Psychopathology Associations

Separate PLS regression models were estimated for the TBI group and the control group. For the TBI group, the first PLS component was statistically significant (*permuted p* = 0.008), with higher scores on the brain latent variable positively associated with greater overall CBCL psychopathology scores (*r* = 0.22). No significant PLS component was identified in the control group, indicating that the observed brain-psychopathology relationship was specific to children with a history of TBI.

Ten of the twenty discriminative neuroimaging features (VIP > 1.0) contributed to the significant TBI-group component (Table 3), spanning all three imaging modalities. The strongest contributors included temporal variance in the right medial orbital sulcus, right calcarine sulcus and left superior parietal lobule, together with gray matter volumes of the left precuneus, central sulcus, inferior occipital gyrus and sulcus, and pallidum (Figure 3).

**Figure 3.**
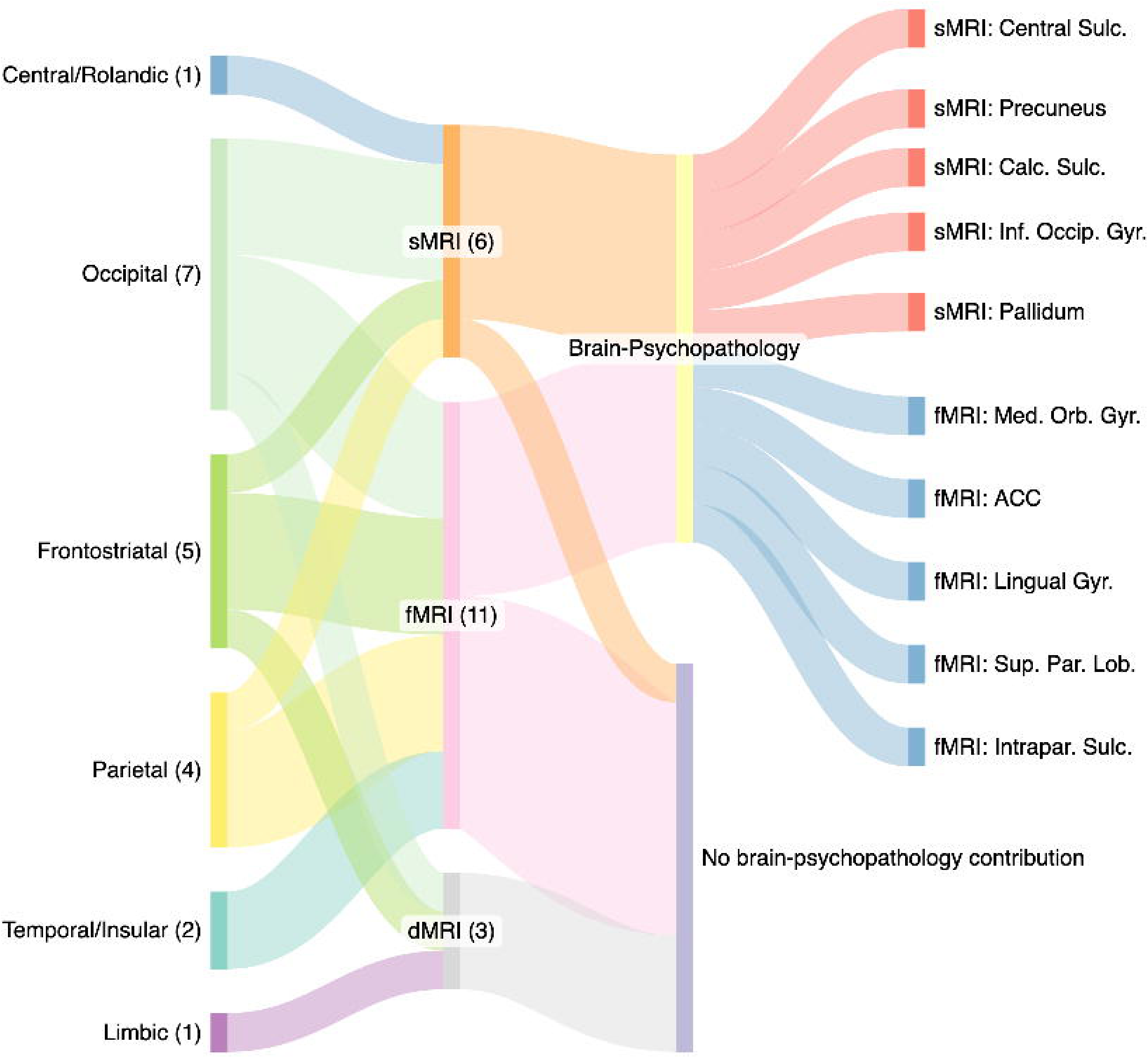
Results of the partial least squares regression between brain imaging features and psychopathology. **(A)** Percentage of variance explained (R²) by each of the first five PLS components (teal bars, and the cumulative variance explained across components (black line). **(B)** Scatter plot of X (brain imaging features) and Y (CBCL psychopathology) scores for Component 1, with a fitted linear regression line (grey) and the corresponding Pearson correlation coefficient (r = 0.274). Each point represents one participant. **(C)** Loadings of individual X variables to Component 1, sorted by magnitude and direction of contribution. Variables with Variable Importance in Projection (VIP) > 1 are shown in teal; those with VIP ≤ 1 are shown in light gray. Variable names are displayed within the plot alongside their corresponding bars. Abbreviations: Anter. = anterior; Circ. = circular; Gyr. = gyrus; L = left; Lat. = lateral; Lob. = lobule; Med. = medial; Mid. = middle; Occip. =occipital; R = right; Sulc. = sulcus/sulci; Sup. = superior; VIP = variable importance in projection.

## DISCUSSION

The present study applied an interpretable ensemble machine learning framework to multimodal MRI data from the ABCD Study to identify neuroimaging markers of pediatric traumatic brain injury (TBI) and examine their relationship with psychopathology. Three principal findings emerged. First, ensemble classifiers achieved modest but reproducible discrimination between children with and without a history of TBI, with similar performance observed in an independent validation sample. Second, the most informative features were distributed across structural MRI, diffusion MRI and resting-state fMRI, indicating that no single imaging modality captured the neural consequences of pediatric TBI in isolation. Third, these discriminative neuroimaging features were associated with psychopathology only in the TBI group, suggesting that the identified multimodal brain signature reflects injury-related neural alterations that contribute to psychiatric vulnerability rather than normative brain-behavior relationships.

Classification accuracy in the range of 61-67% and AUC values of 0.64-0.71 are modest and consistent with those reported by comparable neuroimaging-based machine learning studies classifying participants with TBI from control participants.^22, 51, 52^ The modest ceiling likely reflects the challenge of classifying heterogeneous neurodevelopmental populations from neuroimaging data, where injury severity, time since injury, age at injury, and individual developmental variability all contribute noise.^7, 11^ AUC of the voting, stacking, and bagging ensembles were consistently higher compared to the gradient boosting classifier. This was particularly evident in the independent validation sample, where the boosting classifier achieved accuracy of 0.585. This suggests that probability aggregation across diverse base learners provides more reliable confidence estimates than sequential boosting in this application. The similarity of classification performance between the independent validation sample and the whole sample analysis supports the robustness of the classification framework.

A key strength of the present study is the integration of multimodal neuroimaging. Discriminative features were identified across structural MRI, diffusion MRI and resting-state fMRI, demonstrating that pediatric TBI is characterized by distributed alterations spanning gray matter morphology, white matter microstructure and functional brain organization. This finding is consistent with growing evidence that TBI disrupts multiple interacting neural systems rather than producing isolated focal abnormalities.^53^ Structural MRI captures macroscopic changes in cortical and subcortical anatomy, diffusion MRI provides complementary information regarding white matter integrity, and resting-state fMRI reflects alterations in large-scale functional organization.^7, 12, 13, 15, 16, 18, 21–25, 27, 51, 53–57^ The contribution of all three modalities to classification supports the view that multimodal imaging provides a more comprehensive characterization of TBI than any individual modality alone.^53^

Rather than converging on a single anatomical region, the most informative features were distributed across interconnected frontostriatal, parietal and limbic systems that collectively support executive function, attention, reward processing and emotional regulation. Among these, altered temporal variance within the medial orbitofrontal cortex emerged as one of the strongest contributors to the brain-psychopathology association. The orbitofrontal cortex is particularly susceptible to traumatic injury because of its anatomical location adjacent to the skull base and has consistently been implicated in affective regulation, reward processing and behavioral control.^58, 59^ Structural and functional disruption of orbitofrontal circuitry has been linked to emotional dysregulation following TBI as well as to a range of childhood psychiatric disorders.^60–62^ Its prominence within the present multivariate brain-behavior analysis therefore provides a biologically plausible mechanism through which pediatric TBI may increase vulnerability to subsequent psychopathology.

Additional discriminative features involved the pallidum, superior parietal lobule, precuneus and cingulo-parietal functional connectivity. Although these regions are often discussed individually, they form interconnected networks supporting cognitive control, attentional allocation and self-referential processing.^63^ Alterations across this distributed system are therefore more consistent with widespread network disruption than isolated regional dysfunction. The superior parietal lobule contributes to attentional control and integration of sensory information, while the precuneus occupies a central position within the default mode network and is involved in internally directed cognition.^64–66^ Similarly, the pallidum forms part of frontostriatal circuits implicated in executive function and motivational behavior.^67, 68^ Together, these findings suggest that psychiatric vulnerability following pediatric TBI may arise from disruption of distributed cognitive control networks rather than damage confined to any single cortical or subcortical structure.

White matter microstructure also contributed to classification, particularly reduced fractional anisotropy within the fornix and altered diffusivity within posterior white matter tracts. These findings are consistent with previous diffusion MRI studies demonstrating persistent axonal disruption following pediatric TBI.^16, 19, 22, 23, 57^ The fornix represents a major limbic pathway supporting episodic memory and communication between the hippocampus and wider cortical networks, and is recognized as particularly vulnerable to traumatic axonal injury.^69, 70^ Interestingly, although fornix integrity contributed to classification, it did not emerge as a principal contributor to the brain-psychopathology association identified by PLS. This pattern suggests that some white matter abnormalities may primarily reflect injury burden, whereas alterations involving distributed cortical systems may be more closely related to psychiatric outcomes.

Several discriminative features were also identified within posterior cortical regions, including occipital, temporal, insular and sensorimotor cortices. While these regions individually have diverse functional roles, their collective contribution further supports the diffuse nature of pediatric TBI. The coexistence of occipital volumetric alterations, abnormal visual cortical temporal variance and increased diffusivity within the forceps major is particularly suggestive of coupled structural and functional disruption affecting posterior brain systems. Rather than representing isolated abnormalities, these findings reinforce evidence that pediatric TBI affects distributed brain networks extending beyond regions traditionally associated with cognitive control or emotional processing, often implicated in psychopathology.

One of the most important findings of the present study is that the multivariate brain-psychopathology association identified by PLS was specific to children with TBI. No significant latent variable was identified in the control group despite application of the same analytical approach, indicating that the discriminative neuroimaging features identified by the machine learning models were not simply associated with normal variation in behavioral symptoms.

Instead, the observed brain-behavior relationship appears to reflect injury-related neural reorganization. Although the strength of this association was modest, effect sizes of this magnitude are typical of multivariate brain-behavior analyses performed in large, heterogeneous population cohorts.^39, 71^ Importantly, the combination of permutation-based feature importance with PLS extends the utility of machine learning beyond prediction alone by identifying biologically interpretable neuroimaging features that are specifically linked to psychiatric outcomes following TBI.

Several limitations should be considered. Primarily, the current study is cross-sectional, and although TBI history preceded the brain and behavioral measurements, it is not possible to determine whether the observed brain alterations represent injury-induced changes or reflect pre-existing differences. Longitudinal designs examining brain structure and function before and after TBI are required to establish causal pathways. TBI history in the ABCD Study is defined retrospectively by caregiver report, and exact injury severity, mechanism, and time since injury are not consistently available, limiting subgroup analyses and the characterization of injury dose-response relationships. The modest classification accuracy likely reflects the heterogeneity of the TBI exposure and its neural consequences; future studies with more granular injury characterization may achieve greater discrimination. Additionally, the sample in the current study consisted of a majority of subjects with possible mild TBI. While mild TBI is clinically relevant with regard to heightened subsequent psychopathology risk, it is unclear whether a sample with a higher proportion of more severe TBI cases would yield similar results. Finally, while the multimodal approach used in the current study aims to give a comprehensive picture of TBI-related brain features, these results still reflect local, regional effects. The widespread nature of regions highlighted here supports the systems-wide impact of TBI, which require systems-wide, network-level analytic approaches.

In conclusion, this study demonstrates that interpretable ensemble machine learning applied to multimodal MRI can identify multimodal neuroimaging markers of pediatric TBI within a large population-based cohort. The identified multimodal brain signature, spanning structural, diffusion and functional imaging measures, was specifically associated with psychopathology among children with TBI, supporting a systems-level model of psychiatric vulnerability following childhood brain injury. By combining interpretable machine learning with multivariate brain-behavior modelling, the present work provides a framework for identifying biologically meaningful neuroimaging markers, which may contribute to improved risk stratification and targeted intervention following pediatric TBI.

## Data availability

The de-identified dataset used in this study corresponds to the Adolescent Brain Cognitive Development (ABCD) Study® Curated Data Release 6.0. The data are hosted by the NIH Brain Development Cohorts (NBDC) Data Hub and can be identified using DOI 10.82525/jy7n-g441. Access to these data is restricted to researchers who submit a Data Access Request and sign a Data Use Certification via the NBDC Data Hub website.

## Acknowledgments

Data used in the preparation of this article were obtained from the Adolescent Brain Cognitive Development (ABCD) Study® (https://abcdstudy.org), held in the NIH Brain Development Cohorts (NBDC) Data Hub. This is a multisite, longitudinal study designed to recruit more than 10,000 children aged 9-10 and follow them into early adulthood. The ABCD Study is supported by the National Institutes of Health and additional federal partners under award numbers U01DA041048, U01DA050989, U01DA051016, U01DA041022, U01DA041028, U01DA041042, U01DA041038, U01DA041025, U01DA041086, U01DA041093, U01DA041106, U01DA041117, U01DA041120, U01DA041134, U01DA041148, U01DA041156, U01DA041174, U01DA041159, U01DA041152, U01DA041153, U01DA051037, and U01DA050987. A full list of federal partners is available at the ABCD Study Federal Partners page. A listing of participating sites and a current listing of investigators can be found on the ABCD Study website. ABCD consortium investigators designed and implemented the study and/or provided data but did not necessarily participate in the analysis or writing of this report.

